# Joint Protein Inference Analysis with PyProteinInference Elucidates Biological Understanding of Tandem Mass Spectrometry Data

**DOI:** 10.1101/2024.10.17.618892

**Authors:** Trent B. Hinkle, Corey E. Bakalarski

## Abstract

Selection and application of protein inference algorithms can have a significant impact on the data output from tandem mass spectrometry (MS/MS) experiments, yet its use is often an afterthought in proteomics research due to the inability to apply different inference algorithms in existing analysis systems today. PyProteinInference provides a comprehensive suite of tools to guide researchers through the application of multiple inference algorithms and computation of protein-level, set-based false discovery rates (FDR) from tandem mass spectrometry (MS/MS) data using a unified interface. Here, we describe the software and its application to a K562 whole-cell lysate as well as in a CRAF affinity-purification mass spectrometry experiment to demonstrate its utility in facilitating conclusions about underlying biological mechanisms in proteomic data.

## Introduction

Advances in tandem mass spectrometry (MS/MS) for proteome-level research have driven many novel insights into basic biology and disease^1^. In a typical mass spectrometry-based proteomics experiment, the protein component of interest is first proteolytically digested into constituent peptides for easier analysis by mass spectrometry. This step produces peptides which are more amenable than intact proteins to high-throughput analysis; however, it comes at a cost, severing the relationship between the peptide and its original protein. As this linkage is no longer maintained, it must instead be inferred computationally after the collected mass spectra are matched to their generative peptides using popular algorithms such as MSFragger ^2^, DIA-NN^3^ or Comet^4^. This process of protein inference attempts to reassemble peptides into the list of proteins thought to be present in a sample. While peptide-spectral matching approaches are well established and rely on data provided by ion masses generated during peptide fragmentation, the digestion of proteins into peptides during sample processing leaves behind scant evidence which can be used to determine the original relationships necessary for data interpretation. Thus, despite being a well-described problem for over fifteen years^5^, protein inference is still a challenging area of active research.

Multiple algorithms to perform protein inference have been developed in response to this challenge, including Protein Prophet^6^, Fido^7^, PIA^8^, and Percolator Protein Inference^9^. These tools typically provide mutually-exclusive, widely varying approaches to inference and scoring which are also dataset-dependent. Different combinations of factors, including sequence overlap or others which may not be predictable *a priori*, greatly alter the sensitivity of the results by influencing how peptides are mapped and which proteins are ultimately reported. Lastly, facile usage and comparison of different methods is difficult due to varying requirements for installation, input, configuration, and output, making comparisons between different approaches time consuming and beyond the reach of many biologists. Instead, many studies use a “one size fits all” approach of applying default methods included alongside search algorithms. Nevertheless, relying on such a single approach may fail to accurately represent the dynamics of the underlying proteome^9^ and some experiments may benefit from alternative methods of inference which could help improve biological understanding — if only it were easy to apply and compare them.

The ability to quickly apply and compare multiple protein inference approaches can be critical to understand mechanistic relationships, especially in the context of experiments where sequence similarity of potential biological drivers could obscure results and a thorough consideration of the peptide evidence present for each protein is necessary. Indeed, the multiple protein inference approach proved to be important in improving biological understanding and directing follow-up experiments of a recently reported CRAF APMS experiment^10^.

Surprisingly, no unified protein inference tool exists to conduct such a comparison of leading inference methods. Given the benefits provided by the multiple protein inference approach in the context of this experiment and others, combined with the lack of a comparable public tool, we implemented pyProteinInference, a single, comprehensive software tool to apply multiple protein inference algorithms and associated set-based false discovery rate determinations while providing easy comparison of the results. The software has been implemented as a cross-platform Python package which provides a streamlined, easy-to-use interface to apply multiple protein inference algorithms and can be run on any modern computing system, from individual workstations to high-performance cloud computing environments. The application can be quickly installed using common tools, accepts common input formats, and presents a consistent output, with the aim of assisting the biologist with data interpretation through the use of a multiple protein inference algorithm approach. Here, we introduce the open-source software and present it in the context of both a benchmarking dataset and a real-life experiment.

### Experimental Procedures

#### Implementation of PyProteinInference

PyProteinInference utilizes a standardized workflow (Scheme 1) to quickly apply multiple protein inference methods, contained in an easily-installed Python package. The library supports common input formats including Percolator-generated output^11^. Configuration is achieved via an easy-to-use YAML file, ideal for both interactive use and as part of larger, automated workflows. Once configured, PyProteinInference can perform and compare four different protein inference methods including parsimony^12^, inclusion, exclusion^13^, and peptide-centric^14^. Each protein inference algorithm creates a different peptide to protein mapping thus influencing which proteins are output in final reports: exclusion excludes all non-globally unique peptides from further analysis ensuring that each peptide can only map to one protein; parsimony maps all of the peptides to the minimal set of proteins able to fully explain the results; inclusion assigns peptides to all possible proteins; and peptide centric assigns peptides to all possible proteins and then reduces redundancy by coalescing protein groups based on proteins that map to the exact same set of peptides. In addition to multiple inference assignment methods, differences in protein scoring approaches can also lead to differences in the results^9^. Thus, pyProteinInference implements an exhaustive set of six protein scoring algorithms including common approaches such as multiplicative log-based scoring and best-peptide-per-protein approaches. The package also provides consistent output across all methods, making external comparison and usage of all methods straightforward. More detailed information including all parameter options and more indepth inference algorithm descriptions and can be found in the package documentation. PyProteinInference can be installed using standard Python methods (pip) and run on any modern platform. Source code and package documentation is available at: https://github.com/thinkle12/pyproteininference.

**Scheme 1.**
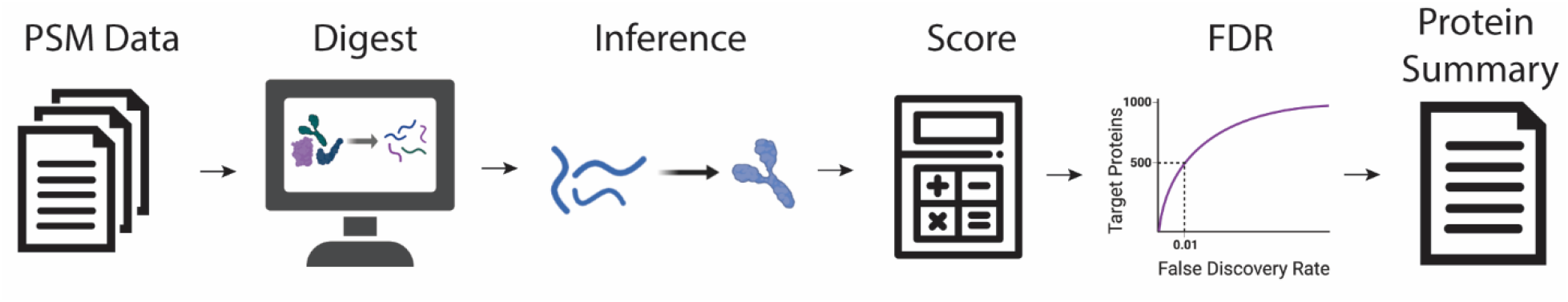
PyProteinInference Workflow.

#### Validation of PyProteinInference algorithms compared to existing methods

To validate the PyProteinInference methods, we created FDR correlation plots of the K562 dataset between PyProteinInference algorithms and corresponding published tools (Figure S1A-C). FDR correlation plots show our implementations to be in agreement with published algorithms. We also compared the overlap of proteins identified in our results against those obtained from previously published tools at a 1% protein FDR (Figure S1D-F). All algorithms in PyProteinInference performed similarly to implementations currently available; the few differences between PyProteinInference and other implementations can be attributed to arbitrary differences in selected proteins due to the stochastic nature of some aspects of the algorithms themselves (i.e., choosing randomly between two assignments of equal weight).

#### Validation of PyProteinInference using K562 whole cell lysate

For algorithm validation of PyProteinInference, we generated a pan-proteome dataset of K562 whole cell lysate using an Orbitrap Fusion Lumos (Thermo Fisher Scientific). First, PSMs were assigned with Comet (v2017.1)^4^ using a concatenated target-decoy database of human proteins (Human SwissProt 2017/08). The Comet search used a peptide mass tolerance of 20 ppm, a fragment bin tolerance of 1.0005 Da, a fragment bin offset of 0.4 Da, and tryptic enzyme specificity with up to 2 missed cleavages. For modifications, a static cysteine carbamidomethyl (+57.02) modification and variable methionine oxidation (+15.99) modification were used. PSM data was then filtered at the peptide level only using Percolator (v3.02.0) with a list of suggested features per PSM^10^ and used as input to PyProteinInference (v1.0.0). PyProteinInference results were generated utilizing three inference options for which comparator software was available (inclusion, exclusion, and parsimony) with posterior error probability as the PSM score and all PSMs from the Percolator results (Table S1). For method validation, inclusion results were obtained from PIA’s (v1.4.5) “report all” method along with parsimony results from its Occam’s Razor algorithm^8^, while exclusion results were generated using Percolator Protein Inference (v3.02.0)^9^ (Table S2). The raw data file and search result files for the K562 dataset described in the text above can be found in the UCSD MassIVE repository (https://doi.org/doi:10.25345/C5KW57N8X) with accession number MSV000089698.

#### Application to CRAF affinity-purification mass spectrometry (APMS) results

To demonstrate the utility of PyProteinInference in data interpretation, we used samples from a previously reported CRAF APMS experiment. Full experimental details, including PSM identification of the raw files and PSM filtering of the kinase domain D468N CRAF dataset, can be found in the supporting publication^10^. After peptide-spectral matching and peptide FDR filtering, all four inference options (inclusion, exclusion, parsimony, and peptide-centric) were performed on the Percolator results and a 4% protein FDR cutoff was used for protein filtering. The output of each inference method coupled with the PSM counts from Percolator filtered to a 1% peptide FDR threshold were then individually run through the SAINT (Significance Analysis of INTeractome) algorithm^15^ to identify potential interacting partners of the kinase domain of the bait protein CRAF with the D468N mutation (Table S3).

## Results and Discussion

### Protein Inference Method Comparison

To showcase the differences between the different protein inference methods and to highlight the importance of using a multiple protein inference approach, we compared the number of identified proteins at a 1% protein false discovery rate (FDR) between each protein inference method integrated into PyProteinInference using the K562 dataset with an upset plot (Figure 1). Examining the individual methods first, we see that peptide centric identifies the most proteins in the K562 dataset (9903) followed by inclusion (6171), parsimony (3099), and finally exclusion (1622) which identifies the fewest proteins at a 1% FDR. Each method identifies measurably different numbers of proteins at an identical FDR. We observed the most restrictive method to be exclusion followed by parsimony, inclusion, and finally peptide centric being the least restrictive method. Additionally, while the different methods have may have considerable similarity in the number of proteins reported overall, they are not identical. For example, the peptide-centric and inclusion methods yield the largest protein numbers yet have differences between their protein sets after intersection (221 proteins). Similarly, while fewer proteins are identified in the parsimony and exclusion methods, 122 proteins differ between the two approaches. This is not unexpected given the different ways in which each method maps peptides to proteins and leads to different significant proteins being identified depending on the protein inference method used, especially in the case of highly similar proteins (such as isoforms). This information should be taken into account when choosing the appropriate inference method for a dataset, which may depend not on maximizing the number of proteins identified but rather on the biological hypothesis being tested; more stringent methods such as exclusion and parsimony can produce higher overall specificity necessary for validating specific mechanistic models while lower sensitivity present in the inclusion and peptide-centric approaches may be more appropriate for screening-based approaches (Figure 1). In any case, given the differences in the number of proteins identified between different inference methods and the measurable set differences between protein inference methods, the benefits of employing a multiple protein inference approach become apparent as the approach maximizes the number of identified proteins while simultaneously providing increased biological context in downstream analyses due to the differences in how each inference method maps peptides to proteins.

**Figure 1.**
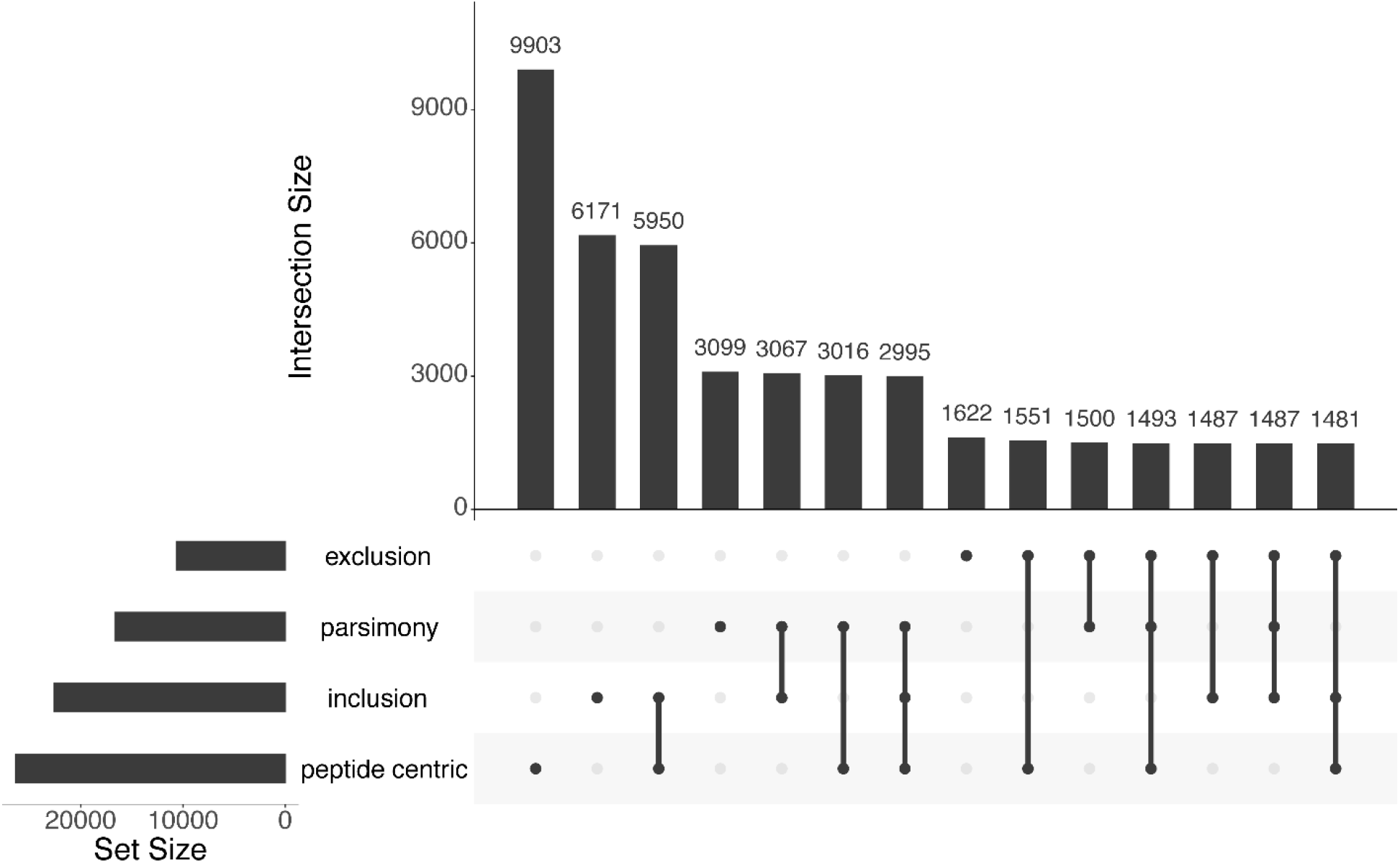
Upset plot showing the overlap of identified proteins at a 1% protein false discovery rate (FDR) from a K562 whole cell lysate dataset. The X axis indicates the individual method or overlapping methods examined while the Y axis indicates the intersection size between methods on the X axis.

### Joint Inference Analysis of affinity-purification mass spectrometry of CRAF

While accurate assignment of peptide-spectral matches to their likely protein counterparts is necessary across most proteome-scale experiments, it is of critical importance when discriminating between closely related interacting partners, where relying on a single inference method may fail to reveal nuances in the data that can support alternate conclusions. In a recent study to determine and validate the genetic conditions necessary for KRAS mutant tumor cell growth^10^, we applied PyProteinInference to generate separate analyses of a D468N mutant CRAF (kinase domain) APMS dataset with alternative protein inference methods. The D468N mutation was hypothesized to induce the loss of kinase activity of CRAF and impact CRAF’s ability to dimerize with ARAF and BRAF. Dimerization events between different RAF isoforms have long been known to play a key role in triggering the RAF/MEK/ERK kinase cascade^16,17^, however, due to close sequence homology of ARAF, BRAF and CRAF in addition to other potential interactive partners MP2K1 and MP2K2, identifying these interactions via mass spectrometry can be difficult. Thus, this experiment provides an interesting opportunity to showcase the utility of the multiple inference method due to this sequence overlap (Figure S2). SAINT^15^ results from each inference method highlight different potential interactors for the kinase domain of the bait CRAF D468N by comparing the CRAF D468N samples against a negative control. Starting with exclusion (Figure 2A), we see that the only potential interacting partners with CRAF are CDC37, MP2K2, and EPIPL; in addition, the bait CRAF does not appear as an interaction in this case. In contrast, the parsimony method indicates CRAF and CLUS in addition to CDC37, MP2K2, and EPIPL (Figure 2B). The inclusion method indicates even more proteins; the same interactions are identified as parsimony in addition to ARAF, BRAF, MP2K2, CUL7 and various isoforms (Figure 2C). Lastly, the peptide-centric approach identifies the same proteins as inclusion, albeit with new protein groups created due to its unique protein grouping strategy (Figure 2D). In the end, applying the same SAINT analysis to datasets utilizing different protein inference algorithms provides for vastly different outcomes and predicted interacting partners for the kinase domain of CRAF with the D468N mutation.

**Figure 2.**
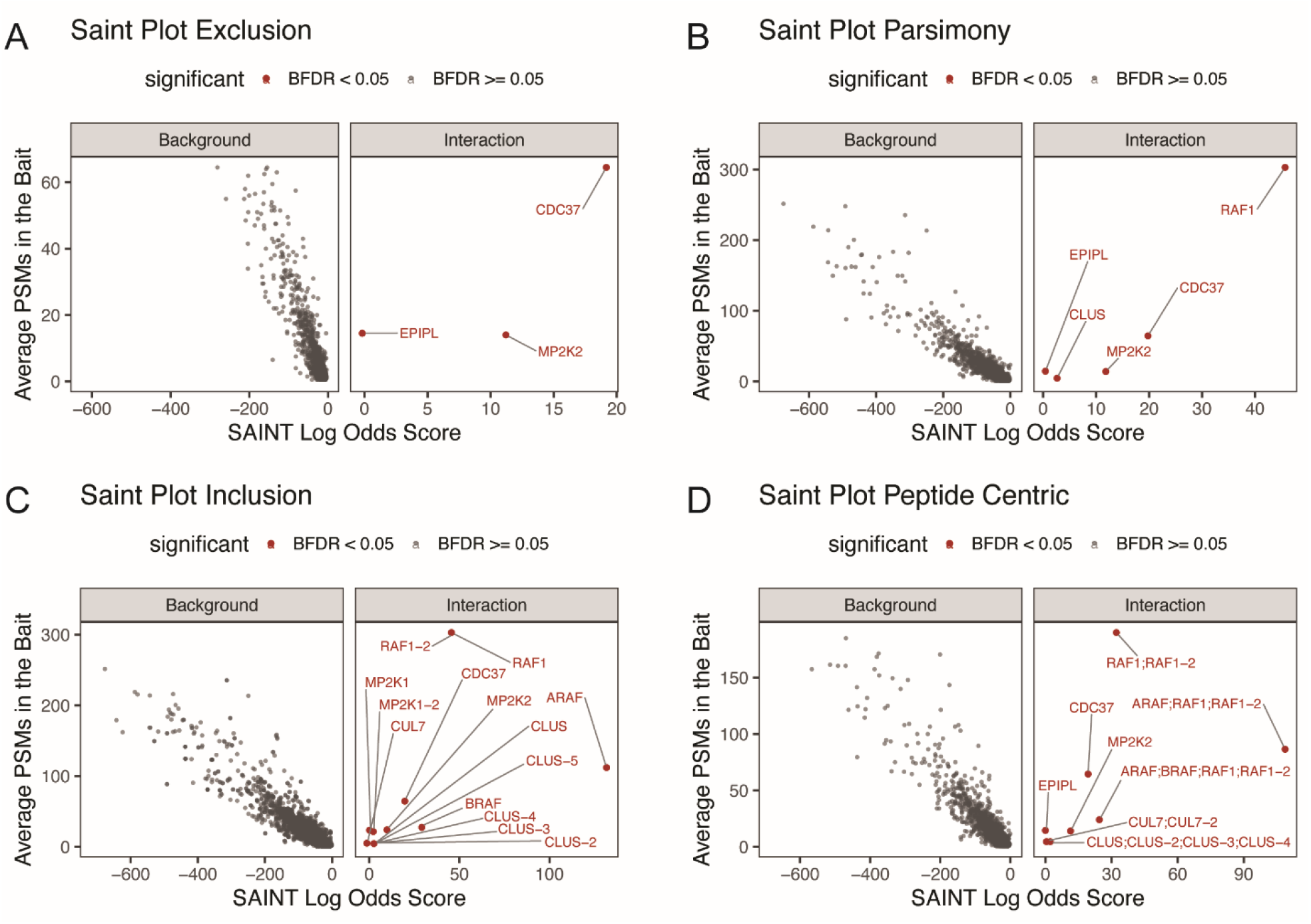
(A-D) SAINT plots that indicate the potential interacting partners with the bait (CRAF) with all four different iterations of the CRAF kinase domain mutation (D468N) APMS dataset, each utilizing a different protein inference method as data input. The axes of the SAINT plots are the SAINT log odds score (the likelihood that a prey is an interacting partner with the bait) against the average number of PSMs in the bait searches. Prey proteins with a Bayesian false discovery rate (BFDR) of less than 0.05% are colored in red and are designated as interacting partners with the bait while proteins with a BFDR greater than 0.05% are designated as background proteins.

Examination of the interacting partners of the kinase domain of CRAF with D468N mutation across all four inference methods pieces together a more accurate story of the potential underlying interactions. The most restrictive approach, the exclusion approach, identifies only MP2K2, CDC37, and EPIPL as distinguishing interactors (as each protein contains at least one peptide unique to it across the entire dataset and each protein only contains unique peptides). Notably, ARAF and BRAF are not identified with the exclusion approach. This seems to indicate that neither ARAF or BRAF have enough globally unique peptides identified in the bait samples to indicate an interaction. Secondly, using the widely-employed parsimony method which generates the minimal list of proteins sufficient to explain all observed peptides, only one additional probable interacting partner is identified, CLUS. Again, ARAF and BRAF were not identified as potential interacting partners despite prior evidence of CRAF:ARAF as well as CRAF:BRAF dimerization^16,17^. Thus, in considering data generated from just this method, it could be concluded that the lack of kinase activity of CRAF (D468N) leads to a loss of CRAF:ARAF and CRAF:BRAF dimerization. However, when the inclusion and peptide centric approaches are considered, ARAF, BRAF, and CUL7 are additionally identified as potential interactors of CRAF. While the majority of the additional peptides are non-distinguishing and also map to the bait CRAF, the interpretation of multiple inference approaches surfaces this fact and reinforces that although there are no unique peptides which unequivocally indicate the presence of ARAF and/or BRAF, this nevertheless does not rule out their presence or activity. Knowledge of the potential interaction of CRAF with ARAF and BRAF depends on having examined the data with these alternative protein inference methods which are infrequently performed. Taken as a whole, support for the absence or presence of ARAF and BRAF is technically ambiguous, and without the further context of parsimony and exclusion, it would be more difficult to come to this conclusion without a deep dive into the data. The utilization of the multiple inference approach possible with PyProteinInference helped dictate follow-up experiments were necessary to accurately determine CRAF kinase domain interacting partners based upon a more detailed examination of the stoichiometric ratios of ARAF and BRAF in the presence of CRAF D468N. Subsequently, these experiments utilized the PIKES^10,18^ method to confirm both CRAF:ARAF and CRAF:BRAF dimerization is occurring in the full length CRAF D468N mutation, with the CRAF:BRAF dimerization being favored. The method also suggested that samples used contain more ARAF than BRAF, which correlated with the average number of PSMs found in bait conditions of peptides mapping to ARAF and BRAF while using the inclusion inference method (Figure 2C, 2D).

Indeed, while from a computational or philosophical perspective algorithms such as parsimony are attractive, they can sometimes be at odds with biological complexity that does not always follow such logical patterns. With these details in mind, the benefits of PyProteinInference become clear, allowing biologists to more accurately explain a biological system with sequence overlap, to be less likely to misinterpret results from one-off protein inference analyses, and to be more prepared for potential follow up experiments given the breadth of context PyProteinInference provides.

## Conclusions

PyProteinInference is a comprehensive software package which implements multiple inference methods for tandem MS/MS data in a single, easy-to-use interface. With the multiple protein inference approach, PyProteinInference provides increased biological context for data interpretation and helps researchers make decisions on potential follow-up experiments by uncovering ambiguity in the data which could support different biological hypotheses. By providing a powerful, integrated tool with a low barrier to entry, PyProteinInference helps researchers maximize understanding of their proteomics datasets and their application to interrogate both biology and disease.

## Supporting Information

- PyProteinInference Comparison to Other Tools (Figure S1)
- CRAF Interacting Partners Sequence Similarity Matrix (Figure S2)
- K562 PyProteinInference Data (Table S1)
- K562 External Protein Inference Data (Table S2)
- CRAF D468N Kinase Domain SAINT Data (Table S3)

## Software Notes

PyProteinInference can be installed using pip. Source code is available at: https://github.com/thinkle12/pyproteininference and package documentation is available at: https://thinkle12.github.io/pyproteininference.

## Acknowledgements

We thank Tommy Cheung for generation of the K562 dataset; figures created with BioRender.com

## References

(1) Yates, J. R. Recent Technical Advances in Proteomics. F1000Research 2019, 8, F1000 Faculty Rev-351. 10.12688/f1000research.16987.1.

(2) Kong, A. T.; Leprevost, F. V.; Avtonomov, D. M.; Mellacheruvu, D.; Nesvizhskii, A. I. MSFragger: Ultrafast and Comprehensive Peptide Identification in Mass Spectrometry–Based Proteomics. Nat. Methods 2017, 14 (5), 513–520. 10.1038/nmeth.4256.

(3) Demichev, V.; Messner, C. B.; Vernardis, S. I.; Lilley, K. S.; Ralser, M. DIA-NN: Neural Networks and Interference Correction Enable Deep Proteome Coverage in High Throughput. Nat Methods 2020, 17 (1), 41–44. 10.1038/s41592-019-0638-x.

(4) Eng, J. K.; Jahan, T. A.; Hoopmann, M. R. Comet: An OpenLJsource MS/MS Sequence Database Search Tool. PROTEOMICS 2013, 13 (1), 22–24. 10.1002/pmic.201200439.

(5) Nesvizhskii, A. I.; Aebersold, R. Interpretation of Shotgun Proteomic Data: The Protein Inference Problem. Mol Cell Proteomics 2005, 4 (10), 1419–1440. 10.1074/mcp.r500012-mcp200.

(6) Nesvizhskii, A. I.; Keller, A.; Kolker, E.; Aebersold, R. A Statistical Model for Identifying Proteins by Tandem Mass Spectrometry. Anal. Chem. 2003, 75 (17), 4646–4658. 10.1021/ac0341261.

(7) Serang, O.; MacCoss, M. J.; Noble, W. S. Efficient Marginalization to Compute Protein Posterior Probabilities from Shotgun Mass Spectrometry Data. J. Proteome Res. 2010, 9 (10), 5346–5357. 10.1021/pr100594k.

(8) Uszkoreit, J.; Maerkens, A.; Perez-Riverol, Y.; Meyer, H. E.; Marcus, K.; Stephan, C.; Kohlbacher, O.; Eisenacher, M. PIA: An Intuitive Protein Inference Engine with a Web-Based User Interface. J. Proteome Res. 2015, 14 (7), 2988–2997. 10.1021/acs.jproteome.5b00121.

(9) The, M.; MacCoss, M. J.; Noble, W. S.; Käll, L. Fast and Accurate Protein False Discovery Rates on Large-Scale Proteomics Data Sets with Percolator 3.0. J. Am. Soc. Mass Spectrom. 2016, 27 (11), 1719–1727. 10.1007/s13361-016-1460-7.

(10) Venkatanarayan, A.; Liang, J.; Yen, I.; Shanahan, F.; Haley, B.; Phu, L.; Verschueren, E.; Hinkle, T. B.; Kan, D.; Segal, E.; Long, J. E.; Lima, T.; Liau, N. P. D.; Sudhamsu, J.; Li, J.; Klijn, C.; Piskol, R.; Junttila, M. R.; Shaw, A. S.; Merchant, M.; Chang, M. T.; Kirkpatrick, D. S.; Malek, S. CRAF Dimerization with ARAF Regulates KRAS-Driven Tumor Growth. Cell Reports 2022, 38 (6), 110351. 10.1016/j.celrep.2022.110351.

(11) Käll, L.; Canterbury, J. D.; Weston, J.; Noble, W. S.; MacCoss, M. J. Semi-Supervised Learning for Peptide Identification from Shotgun Proteomics Datasets. Nat. Methods 2007, 4 (11), 923–925. 10.1038/nmeth1113.

(12) Zhang, B.; Chambers, M. C.; Tabb, D. L. Proteomic Parsimony through Bipartite Graph Analysis Improves Accuracy and Transparency. J. Proteome Res. 2007, 6 (9), 3549–3557. 10.1021/pr070230d.

(13) The, M.; Edfors, F.; Perez-Riverol, Y.; Payne, S. H.; Hoopmann, M. R.; Palmblad, M.; ForsstroLJm, B.; KaLJll, L. A Protein Standard That Emulates Homology for the Characterization of Protein Inference Algorithms. J. Proteome Res. 2018, 17 (5), 1879–1886. 10.1021/acs.jproteome.7b00899.

(14) Meyer-Arendt, K.; Old, W. M.; Houel, S.; Renganathan, K.; Eichelberger, B.; Resing, K. A.; Ahn, N. G. IsoformResolver: A Peptide-Centric Algorithm for Protein Inference. J Proteome Res 2011, 10 (7), 3060–3075. 10.1021/pr200039p.

(15) Choi, H.; Larsen, B.; Lin, Z.-Y.; Breitkreutz, A.; Mellacheruvu, D.; Fermin, D.; Qin, Z. S.; Tyers, M.; Gingras, A.-C.; Nesvizhskii, A. I. SAINT: Probabilistic Scoring of Affinity Purification–Mass Spectrometry Data. Nat Methods 2011, 8 (1), 70–73. 10.1038/nmeth.1541.

(16) Farrar, M. A.; Alberola-lla, J.; Perlmutter, R. M. Activation of the Raf-1 Kinase Cascade by Coumermycin-Induced Dimerization. Nature 1996, 383 (6596), 178–181. 10.1038/383178a0.

(17) Yuan, J.; Ng, W. H.; Lam, P. Y. P.; Wang, Y.; Xia, H.; Yap, J.; Guan, S. P.; Lee, A. S. G.; Wang, M.; Baccarini, M.; Hu, J. The Dimer-Dependent Catalytic Activity of RAF Family Kinases Is Revealed through Characterizing Their Oncogenic Mutants. Oncogene 2018, 37 (43), 5719–5734. 10.1038/s41388-018-0365-2.

(18) Reichermeier, K. M.; Straube, R.; Reitsma, J. M.; Sweredoski, M. J.; Rose, C. M.; Moradian, A.; Besten, W. den; Hinkle, T.; Verschueren, E.; Petzold, G.; Thomä, N. H.; Wertz, I. E.; Deshaies, R. J.; Kirkpatrick, D. S. PIKES Analysis Reveals Response to Degraders and Key Regulatory Mechanisms of the CRL4 Network. Mol. Cell 2020, 77 (5), 1092-1106.e9. 10.1016/j.molcel.2019.12.013.

