## Supplementary figures and images for "Joint Protein Inference Analysis with PyProteinInference Elucidates Biological Understanding of Tandem Mass Spectrometry Data"

### Supplemental Figures

#### Supplemental Figure 1

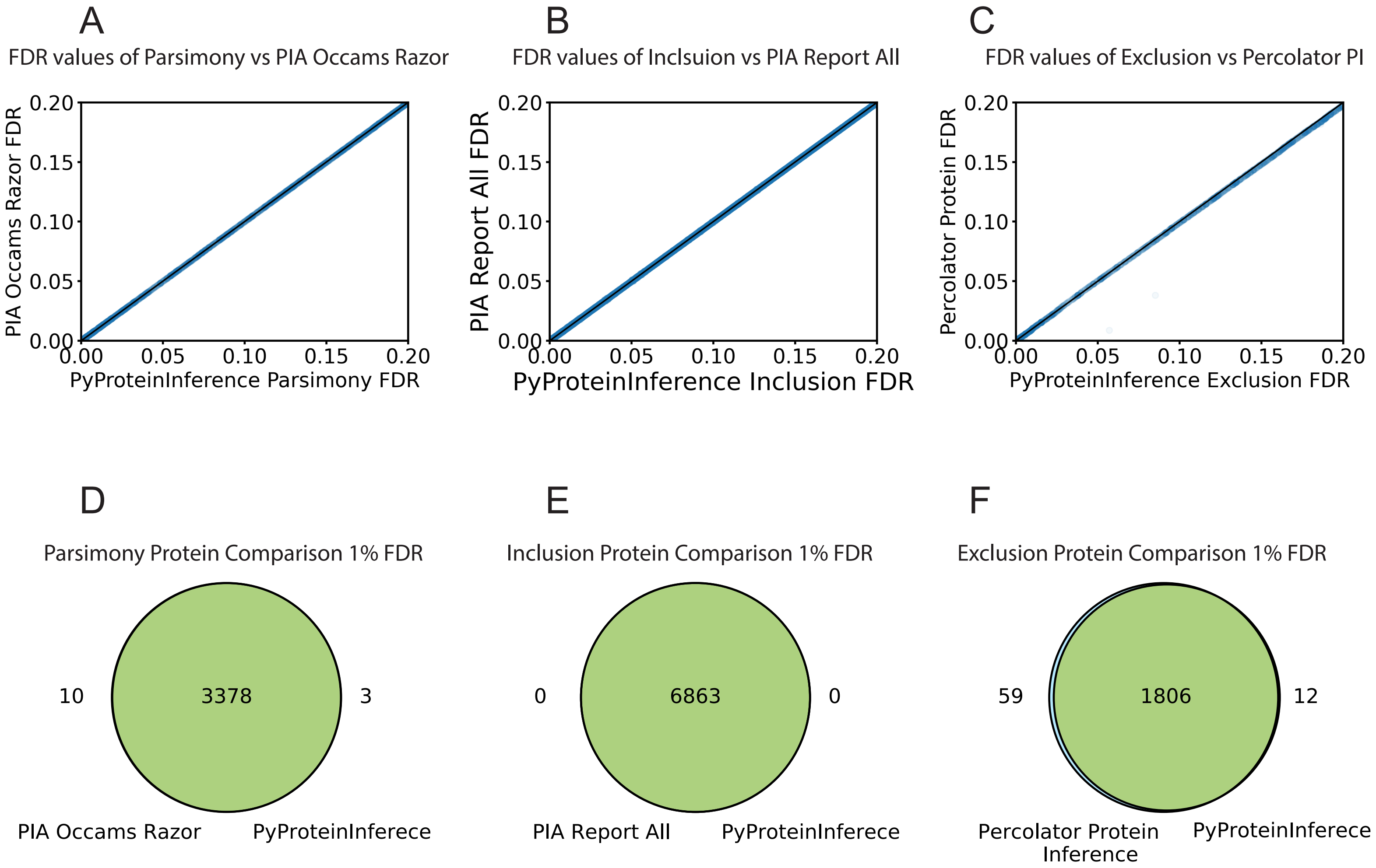

Supplemental Figure 2

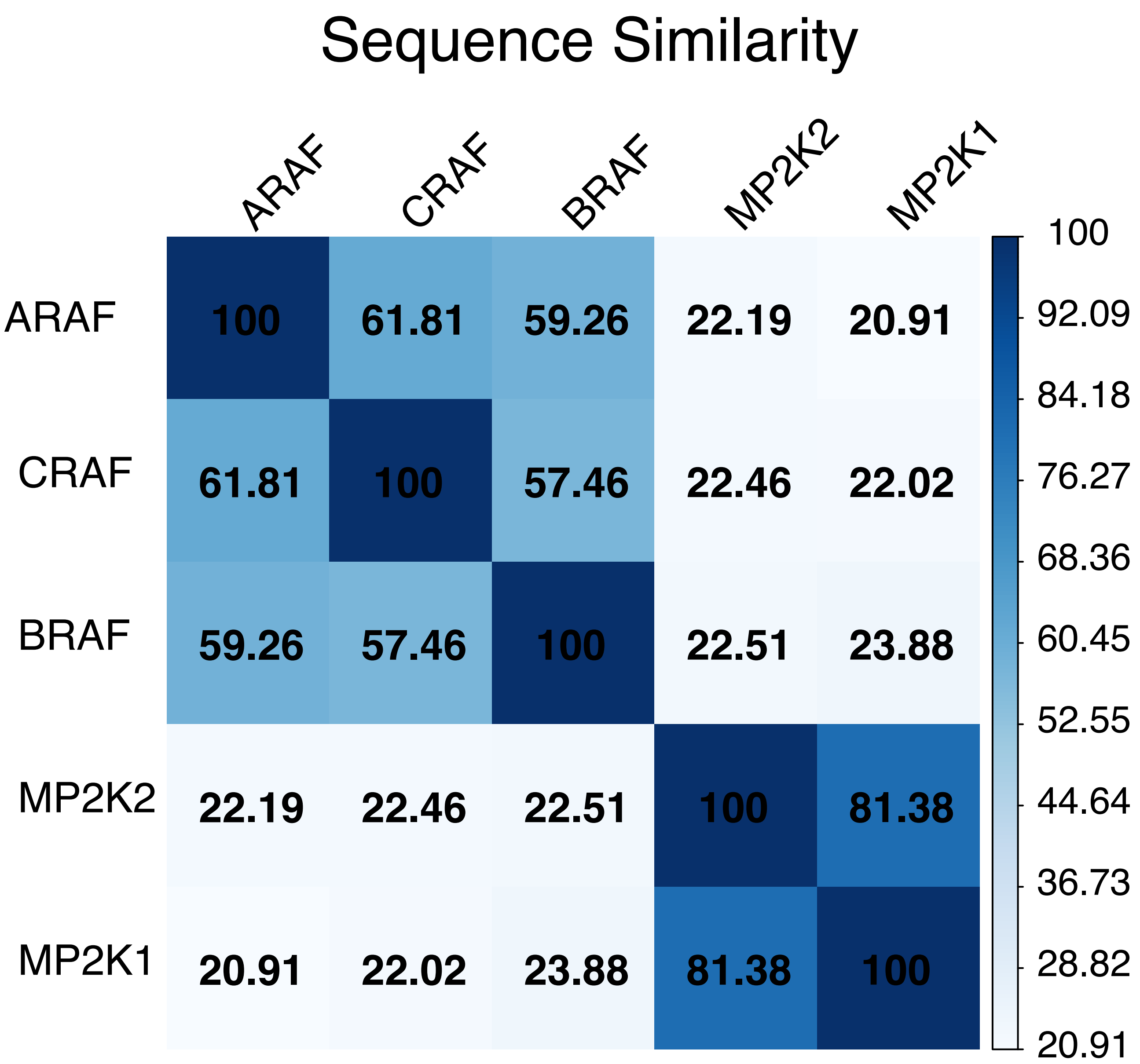
